# Division of labor increases with colony size, regardless of group composition, in the social spider *Stegodyphus dumicola*

**DOI:** 10.1101/652677

**Authors:** Colin M. Wright, James L. L. Lichtenstein, C. Tate Holbrook, Justin Pretorius, Noa Pinter-Wollman, Jonathan N. Pruitt

## Abstract

Division of labor (DOL) is a pattern of work organization where individual group members specialize on different tasks. DOL is argued to have been instrumental for the success of eusocial insects, where it scales positively with group size both within and across species. Here we evaluate whether DOL scales positively with group size in a society of cooperative breeders (social spiders) and whether this pattern is impacted by the behavioral composition of the group. To do this we engineered experimental colonies of contrasting group sizes and behavioral compositions and tracked individuals participation in two colony maintenance tasks: prey capture and web construction. As with some eusocial insects, we found that larger groups exhibited DOL metrics up to 10-times greater than smaller groups, conveying that individuals specialize on particular tasks more in larger colonies. This scalar relationship did not differ by a groups behavioral composition, though groups composed of only bold spiders exhibited reduced DOL relative to all-shy or mixed groups. We also found that per capita participation in prey capture, but not web construction, decreased as a function of group size. This suggests that individuals in larger groups may save energy by reducing their involvement in some tasks. Together, our results convey that similar scalar relationships between DOL and group size can emerge both inside and outside the eusocial insects. Thus, theory developed for understanding DOL in eusocial societies may inform our understanding of group function in a larger swath of animal social diversity than is broadly appreciated.

**Significance Statement:** Division of labor (DOL) has been a major area of research in the eusocial insects for decades, and is argues to underlie their ecological success. Only recently have other social arthropods, such as social spiders, been considered for studies concerning DOL. Given their smaller colony sizes, and absence of morphological castes, DOL was not thought to be an important facet of spider societies. However, we found that spider societies do indeed exhibit high degrees of DOL that is positively correlated to colony size, as seen in many eusocial insects. These findings suggest that the scalar relationship between group size and social organization seen in social insects is likely generalizable to a larger diversity of social taxa, and that cooperative breeders can show levels of division of labor equaling or exceeding those of eusocial systems evaluated to date.

## Introduction

The concept of Division of labor (DOL) was first articulated by the economist Adam Smith in his book *The Wealth of Nations* (1776). In it, he described a process whereby workers can increase group productivity and output by assigning different jobs to different individuals. The increase in productivity and efficiency produced by DOL is attributed to several factors: (1) time saved by the elimination or reduction of task switching, (2) reduced task learning costs, and (3) increased task efficiency associated with practiced motor learning and dexterity. In many social taxa, including humans, individuals often differ innately in their aptitudes for different tasks. Thus, DOL and within-group behavioral variation can allow workforces to adaptively align individuals with contrasting aptitudes with the tasks that they each perform best (Oster and Wilson 1978). Although DOL was originally formulated in the context of the human factory and assembly-line workers, it has since been recognized as a ubiquitous phenomenon in biological systems from cells to social insect colonies (Szathmary and Smith 1995). Many examples have been identified across the animal kingdom, where DOL is defined as a pattern of non-random association between individual worker identities and the tasks that they perform (Michener 1974; Rypstra 1993; Duarte et al. 2011a; Jeanson 2013; Jeanne 2016).

While research on division of labor has traditionally focused on eusocial insects, which exhibit reproductive DOL between queen and worker castes (Oster 1978), other animal societies further divide non-reproductive labor among workers or group members. Interindividual variation in task performance can be associated with differences in morphology (e.g., majors vs. minors) (Wilson 1980; Holldobler and Wilson 1990; Kay and Rissing 2005), age (temporal polyethism) (Wilson 1971; Seeley 1982), innate response thresholds (Bonabeau et al. 1996; Bonabeau et al. 1998; Beshers 2001; Jeanson 2013), reproductive state (social spiders: (Junghanns et al. 2017), or consistent individual differences in behavioral tendencies (Dornhaus 2008; Holbrook et al. 2014; Wright et al. 2014). This last element of individual variation can arise from a variety of mechanisms including individual differences in genotype, gene expression, hormonal changes, and learning/experience (Duarte et al. 2011b). In some cases, such differences can cause individuals to perform better at some tasks than others (Oster 1978; Holldobler and Wilson 1990).

DOL, however, is not a simple, static trait. Rather, it is a dynamic emergent property of a group. Within a species, for instance, DOL tends to increase with colony size (Holbrook et al. 2011). This *social scaling effect* is consistent with a general principal known as the *size-complexity rule* (Bonner 2004) that applies to many biological entities, including human societies. This rule states that, in general, larger collective entities have higher DOL than smaller entities, and these “entities” can refer to individual organisms composed of various types of cells or cooperative societies composed of various types of individuals. The scaling of DOL in functionally integrated societies could be an evolved response that allows them to overcome logistical challenges associated with increasing group size, and thus, helps to increase performance efficiency. Alternatively, the positive relationship between DOL and group size could be an emergent consequence of social dynamics, as predicted by self-organizational models (Gautrais et al. 2002; Merkle and Middendorf 2004; Jeanson et al. 2007; Ulrich et al. 2018). This latter hypothesis has received some support from empirical tests in forced associations of normally solitary ant foundresses (Jeanson and Fewell 2008) and sweat bees (Holbrook et al. 2013).

The DOL index developed by Gorelick et al. (2004) and Gorelick and Bertram (2007) provides researchers with a rigorous and generalizable method to measure DOL and make comparisons across social groups and taxa. In short, their approach quantifies the degree to which individuals specialize on a subset of tasks instead of dividing their time evenly across all tasks, and the degree to which different individuals specialize on different tasks. This allows for both intra- and interspecific comparisons of division of labor to be made, and allows researchers to detect DOL in species that lack morphological worker subcastes or spatially-linked DOL. The transitionally social spider *Anelosimus studiosus*, for instance, exhibits one of two behavioral types (docile or aggressive) that can occur either solitarily or in groups (Riechert and Jones 2008). Individual personality type as well as the behavioral composition of *A. studiosus* colonies have been shown to influence participation, behavioral plasticity, and individual proficiency in tasks (Pruitt et al. 2012; Pruitt 2013; Holbrook et al. 2014; Wright et al. 2014). Furthermore, *A. studiosus* exhibits levels of DOL (Holbrook et al. 2014) on par with those of bumblebees (*Bombus impatiens*) (Jandt et al. 2009) and communal sweat bees (*Lasioglossum hemichalceum*) (Jeanson et al. 2005), but lower than values observed in eusocial species such as harvester ants (*Pogonomyrmex californicus*) (Jeanson and Fewell 2008) or rock ants (*Temnothorax albipennis*) (Dornhaus et al. 2009). Results like these convey that convergent patterns of social organization can emerge across a spectrum of social systems and taxa. This, in turn, suggests a greater taxonomic zone of application for theories developed within the social insect literature.

The African desert social spider, *Stegodyphus dumicola*, is an excellent model system to investigate personality-linked DOL. Individuals within *S. dumicola* colonies exhibit high levels of “personality” variation in boldness, and a colony’s personality composition has been shown to be a significant predictor of collective outcomes such as collective foraging aggressiveness (Pruitt and Keiser 2014), colony defense (Wright et al. 2016), web repair (Keiser et al. 2016), and bacterial transmission rates (Keiser et al. 2017). Previous studies have demonstrated that *S. dumicola* colony members exhibit some degree of personality-linked task differentiation in colony defense and foraging tasks (Wright et al. 2015; Wright et al. 2016). However, the extent to which these animals divide labor across multiple tasks in colonies of various sizes and compositions is unknown.

Here we query whether *S. dumicola* colonies divide labor based on individual differences in personality, and how this relationship changes as a function of group size. We further examine how *per capita* participation in colony maintenance tasks scales as a function of group size, as the work needed to maintain a colony relative to workforce size likely varies (Jeanson et al. 2007). Specifically, we ask the following questions: (1) To what degree does *S. dumicola* exhibit DOL across prey capture and web building? (2) How does personality composition affect the expression of DOL? (3) Does DOL scale with colony size? And, if so, (4) does this scalar relationship change based on a group’s personality composition? It is important to note that many factors, such as developmental stage, time of day, and other environmental factors may influence task participation and division of labor. In prior *S. dumicola* studies, we showed that individuals’ personalities are linked with their probability of performing various tasks within the group (Wright et al. 2015; Wright et al. 2016), as well as individuals’ tendencies to respond to the behaviors of others (Pruitt et al. 2017). We therefore chose to explicitly test for the effects of personality composition and colony size on DOL, while holding as many other variables constant as possible (e.g., using only subadults, testing at the same time each day, holding conditions constant in greenhouse conditions). We predicted that greater within-group behavioral variation will enhance DOL because of personality-linked task differentiation, and that this effect will be most pronounced in larger groups.

## Methods

### Data Collection Overview

The data herein were collected as part of a senior level animal behavior class at the University of California at Santa Barbara. Several precautions were taken to ensure publication worthy data. First, all students received a four-week training period prior to data collection. Students were trained on proper handling of spiders, conducting collective prey capture assays, color IDing individuals, and identifying web components (capture web vs. retreat). Second, data collection was monitored closely at all times by three senior observers: the PI, a TA, and a undergraduate TA. Third, all data were collected by groups of students (3-4 students/group) and no students were ever permitted to collected data independently or without a senior observer present. Fourth, the colonies assigned to each student group were alternated across days to prevent subtle differences in data collection practices from biasing the outcome of particular treatment groups. Finally, the student observers were blind to the behavioral compositions of the focal colonies. However, observers were not blind with regard to colony size, because larger colonies are conspicuously larger.

### Colony creation and behavioral assessment

Eleven colonies of subadult *S. dumicola* were collected along fences around Upington in the Northern Cape of South Africa, and moved to the lab at the University of California at Santa Barbara in the spring of 2017. All colonies were fed to satiation with 6-week-old domestic crickets as soon as they arrived at the lab, and our experiments began the following week. Colonies were likewise fed an ad libitum meal of domestic crickets once weekly during the course of our studies. Source colonies were sorted and each spider was individually isolated from its nest mates in small 30ml plastic condiment containers. Given that spiders were subadults, we were not able to determine the sex of individual spiders, though natural populations exhibit a 10:1 female to male sex ratio. We then measured each spider’s boldness.

Boldness was evaluated by administering two gentle puffs of air to spiders’ anterior prosomas using a rubber squeeze-bulb. These air puffs are meant to resemble an attack from an avian predator, causing the spider to cease activity and pull its legs in against its body in a “huddle” position (Riechert and Hedrick 1990; Lohrey et al. 2009; Pruitt et al. 2013). For most web-building spiders, sudden directional movements of air simulate like aerial predators (Foelix 2011). The latency to unhuddle and move one whole body length following these air puffs is our measure of boldness. Boldness is defined as the propensity of an individual to engage in risky behavior (Sloan Wilson et al. 1994), and resuming activity quickly after interacting with a possible predator (air puffs) is deemed risky or “bold” behavior. Bold and shy individuals are defined as those spiders having latencies falling between 0-299 and 300-600 seconds respectively (Keiser et al. 2014). Boldness is highly repeatable at the individual level (repeatability = 0.63) in this species (Keiser et al. 2014). These boldness scores are then subtracted from 600 (the maximum value) so that higher numbers reflect greater boldness scores. After boldness assays, we measured each spider’s mass and prosoma width to obtain estimates of body size and condition. Body condition was estimated using the residuals of a linear regression of mass on prosoma width, such that spiders that are relatively heavy for their body size are deemed in better condition (Jakob et al. 1996).

After assessing spider morphometry and behavior, we assembled experimental colonies (*N* = 33) composed of 100% bold individuals (*N* = 7), 100% shy individuals (*N* = 16), and a 50/50 mixture of bold and shy individuals (*N* = 10). Colonies ranged in size from 6 to 40 individuals based on the number of spiders of varying personality that we could acquire from our source colonies. Colonies were constructed by sub-setting individuals from a given source colony, thus experimental colonies were always composed of individuals with natural levels of relatedness and familiarity (Seibt and Wickler 1988). Both of these factors impact colony performance in several species of *Stegodyphus* (Ruch et al. 2009) including *S. dumicola* (Laskowski et al. 2016). We painted a haphazard subset of spiders per colony (6-24 individuals/colony, each with a unique color combination) using a Sharpie Paint Pen, and then established colonies on *Acacia reficens* plants kept in a greenhouse, and gave them 24 hours to construct retreats and capture webs. 100% of colonies produced capture webs during this time.

We measured each colony’s responsiveness to prey by vibrating a small (1cm^2^) piece of white paper placed in the center of the capture webs with a handheld vibrator. This causes the paper to flutter about in a manner reminiscent of a struggling winged insect, and spiders respond to this stimulus by emerging from their retreat to attack the vibrating paper. Once we began vibrating the paper, we recorded the unique color ID of each spider participating in the attack. The vibrational stimulus occurred for a maximum of 5 minutes, and all colonies responded within this time frame for each trial. An “attack” was deemed to have occurred when spiders responded to the vibrational stimulus and made contact with the paper. The number of participants was measured as the number of spiders on the capture web actively approaching the paper at the moment the first individual made contact with it. These assays were performed twice daily for five consecutive days, followed by a two-day rest period. We then performed the assay twice daily for another five consecutive days (20 total observations per colony). Following these prey capture assays, we observed each colony during the night between 8:00-9:00pm for four nights (2 observations per week, on Monday and Friday) and recorded the total number of spiders and color ID of each spider that we observed constructing new webbing. None of the marked individuals molted during the experiment.

### Ethics Statement

At the end of the experiment colonies were maintained in the greenhouse environment until all the spiders perished, seemingly of old age, as their longevity in laboratory exceeds those of *S. dumicola* in the wild. Although invertebrates are not subject to Institutional Animal Care and Use Committee (IACUC) regulation in the United States, care was taken not to unnecessarily stress the animals and their longevity resembled those of individuals in their native habitat.

### Data analysis: division of labor assessment

Division of labor was quantified following Gorelick et al. (2004). For each colony, across all observations, we generated an individual × task data matrix where each cell contained the frequency with which we observed a specific individual participating in a specific task: prey capture and web construction. We normalized the matrix so that the sum of all entries equaled 1. From the normalized matrix, we calculated Shannon’s index, or marginal entropy, over individuals (*H*indiv) and over tasks (*H*tasks):

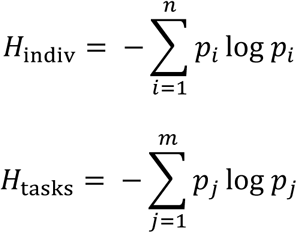

where *pi* is the probability that the *i*th individual performed any task and *pj* is the probability that any individual performed the *j*th task. We then calculated mutual entropy between individuals and tasks (*I*_indiv,tasks_), given by

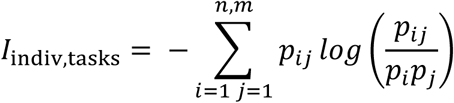

where *pij* is the joint probability that the *i*th individual performed the *j*th task. Finally, the DOL indices are defined as follows.

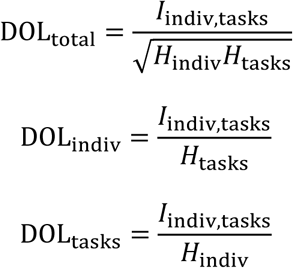

In our study, we chose to focus on DOL_indiv_ for our measure of division of labor, as this metric is insensitive to differences in group size, and can therefore be used to test for actual group size effects on division of labor. The full derivation of DOL indices can be found elsewhere (Gorelick et al. 2004), but note that the definitions of DOLindiv and DOLtasks were accidentally switched in the original manuscript.

### Data analysis: group-level effects

We tested for the effects of colony behavioral composition (100% shy, 100% bold, and mixed) and colony size on DOL using GLMM with a normal distribution and identity-link function. Our predictor variables were group behavioral composition (bold, shy, and mixed), colony size, and a composition × colony size interaction term. Colony ID was nested within source colony ID, and both were included a random effects in our analysis. The DOL_indiv_ metric for each colony was our response variable. A significant interaction term between colony size and composition was then used to test whether the scalar relationship between colony size and DOL varied based on a colony’s behavioral composition.

We constructed three LMM to evaluate whether colonies’ degree of DOL were associated with the average mass of group members, the average prosoma width of group members, or the average body condition of group members. A separate model was run for each variable, and source colony ID was included as a random effect in each model. Body condition was evaluated as the residuals of a mass on prosoma width linear regression, where heavier bodied spiders relative to their size are deemed in better condition (Jakob et al. 1996).

### Data analysis: individual-level effects

To evaluate whether individual’s per capita task participation varied as a function of group size, we calculated the portion of observation periods were we observed each marked individual engaged in a task: web construction or prey capture. We then averaged these values within each colony to obtain a single colony level metric for each task type. We then constructed two models, one for each task type, to observe whether per capita involvement in each task decreased with increasing group size. We included colony size as a predictor variable. We used a normally distributed GLMM with identity-link function for this analysis. Source colony ID was included as a random effect in this analysis.

## Results

Our experimental compositions exhibited the following mean DOL_indiv_ values: bold = 0.265, 95% CI [0.203, 0.327], SE [0.0315]; shy = 0.406, 95% CI [0.288, 0.524], SE [0.0602]; and mixed = 0.364, 95% CI [0.257, 0.471], SE [0.0546]. The total average DOL_indiv_ across all compositions was 0.364, 95% CI [0.295, 0.432], SE [0.0348].

For colony-level effects, we found that colony DOL scaled positively with colony size (colony size: t(1) = 2.58, *p* = 0.015, SE = 0.0054), but this relationship did not differ among colonies of different behavioral compositions (composition × colony size: t(2) = 1.4, *p* = 0.38, SE = 0.0093)(Figure 1). We found a significant main effect of behavioral composition as well (composition: t(2) = 1.31, *p* = 0.016, SE = 0.052), where colonies composed of all bold individuals exhibit lower DOL than colonies composed of all shy or a mixture of bold and shy individuals, which were statistically indistinguishable from each other.

**Figure 1:**
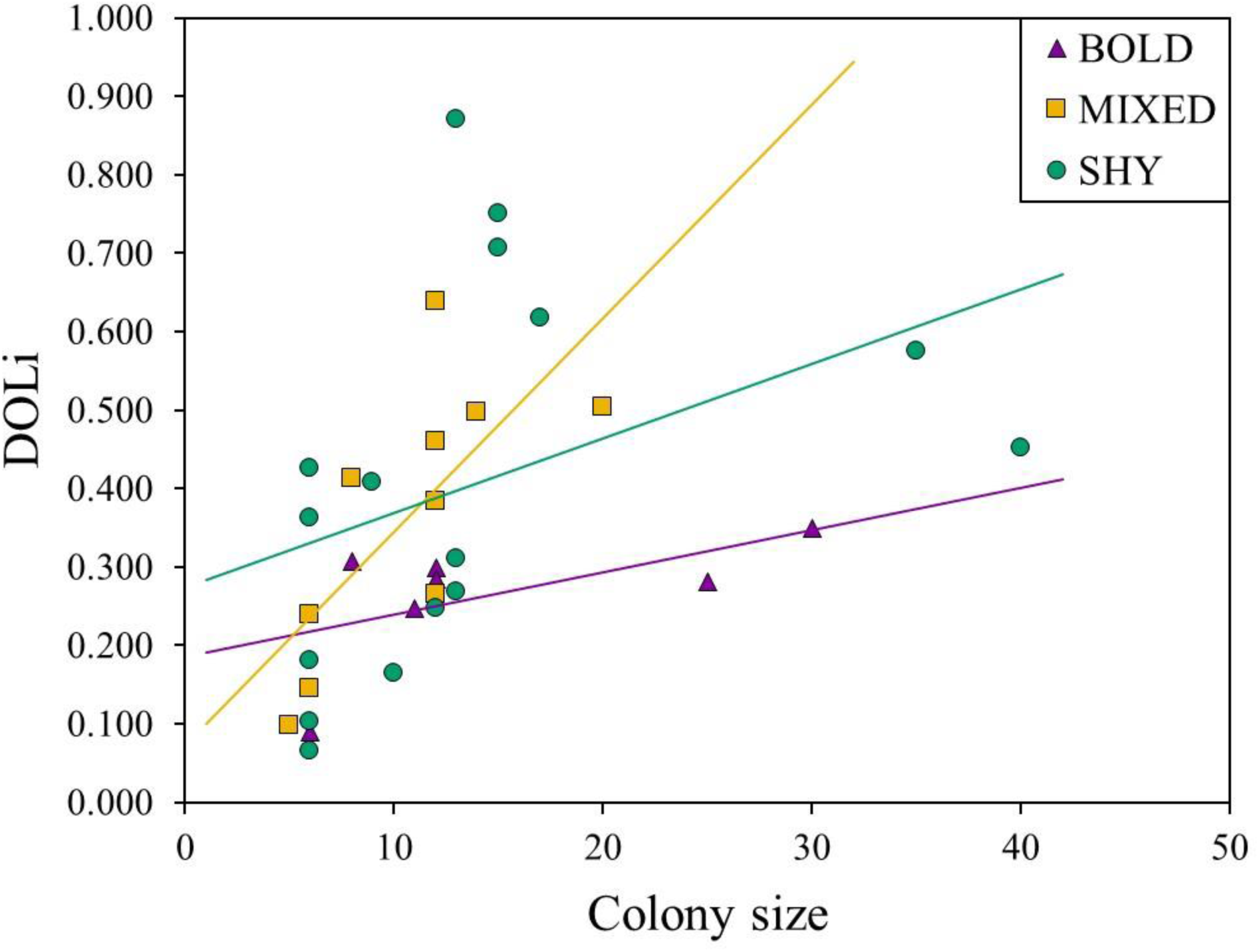
Division of labor (DOL_indiv_) of three behavioral compositions (bold, shy, and mixed) as a function of colony size. Each dot represents the DOL metric calculated for each colony.

We found that colonies’ DOL were unrelated to colony member mass (mass: t(1) = 0.74, *p* = 0.47, SE = 3.08), prosoma width (prosoma width: t(1) = −0.12, *p* = 0.91, SE = 0.13), or body condition (body condition: t(1) = 2.00, *p* = 0.16, SE = 5.81).

We found no relationship between colony size and per capita participation in web building (colony size: t(1) = −0.04, *p* = 0.97, SE = 0.0015), but found a negative relationship between colony size and per capita participation in prey capture (colony size: t(1) = −2.16, *p* = 0.0392, SE = 0.0014) (Figure 2). Source colony had no significant effect on any colony- or individual-level response variable.

**Figure 2:**
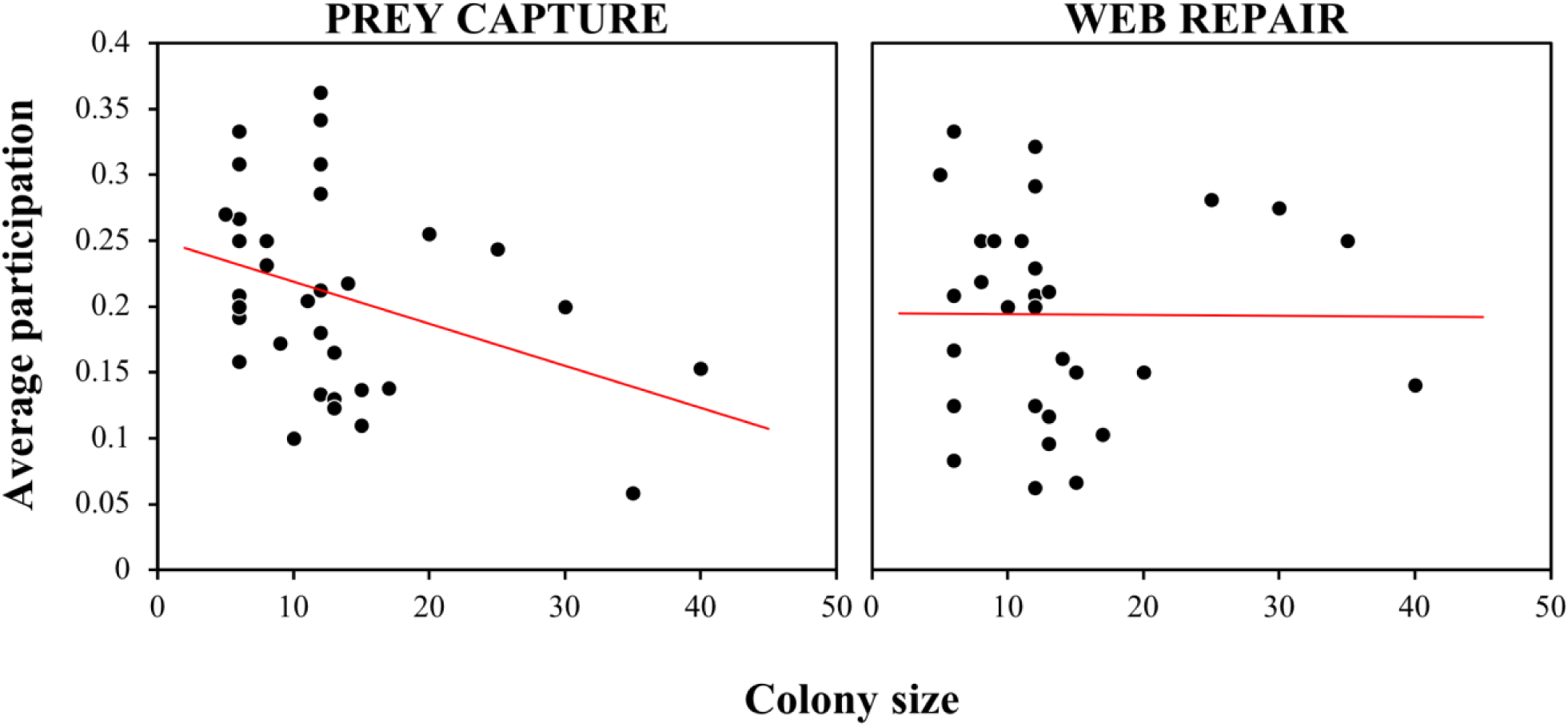
The average proportion of spiders participating in prey capture (left) and web repair (right) as a function of colony size.

## Discussion

Here we show that cooperatively breeding social spiders can exhibit DOL metrics approximating those of some eusocial insects, even those in large colonies (e.g., harvester ants, rock ants). DOL also increases positively with group size in these spiders, as observed in other taxa, but this scalar relationship is unrelated to group composition. DOL was higher, overall, in all-shy and mixed colonies than in all-bold colonies, regardless of colony size. Thus, as in other taxa, DOL appears to be an important organization phenomenon governing patterns of work in *S. dumicola –* although its adaptive significance remains unknown. At the individual level, we found that *per capita* involvement in one colony maintenance task (prey capture) but not another (web construction) decreased with increasing group size. This finding indicates that individuals in larger groups may benefit from decreased energy expenditure and reduced risk associated with some colony maintenance tasks.

At the group level, we observed that groups composed of all bold individuals exhibited lower DOL metrics than other compositions. The tendency for bold individuals to participate more in both prey capture (Wright et al. 2015) and web repair (Keiser et al. 2016) may help to explain this observation. If bold individuals are more likely to execute both tasks, then individuals in all-bold groups may engage in both tasks more, and thus, no individual specialization (engaging in task A but not task B, and *vice versa*) emerges. In contrast, all-shy groups exhibited some of the highest DOL metrics (Figure 1). This finding suggests that pre-existing within-group behavioral variation alone is insufficient to forecast the degree of DOL that emerges within colonies. If it were, then we would have predicted the greatest DOL in groups composed of a mixture of behavioral phenotypes (Holbrook et al. 2014). This discrepancy may be in part due to the limited scope of observed tasks, notably the exclusion of inside-retreat tasks such as brood care, which may be disproportionately performed by shy individuals. However, bold individuals may be less socially responsive than their shy counterparts, which may decrease DOL in social rodents (Koolhaas et al. 1999), fish (Jolles et al. 2015) but see (Trompf and Brown 2014), birds (Kurvers et al. 2009; Kurvers et al. 2010), the transitionally social spider *A. studiosus* (Holbrook et al. 2014), and even in *S. dumicola* (Pinter-Wollman et al. 2017). If shy *S. dumicola* are indeed more socially responsive than their bolder counterparts, as argued elsewhere (Pinter-Wollman et al. 2017; Pruitt et al. 2017), then it follows that shy individuals might be better able to sense the absence of bold individuals (or elevated task-related stimuli in their absence) and begin to perform the tasks typically executed by them (as seen in *A. studiosus*: Holbrook et al. 2014). Regardless of the precise mechanism involved, in aggregate, it appears that the presence of shy individuals is an important causal driver of DOL in *S. dumicola*.

As in many insects (Jeanne 1986; Gordon 1989; Thomas and Elgar 2003; Dornhaus et al. 2009; Holbrook et al. 2011), but see Dornhaus et al. (2009), DOL scales positively with group size in *S. dumicola*. In insects, DOL is thought to enhance colony performance and allow for the evolution of larger and more complex colonies (Oster and Wilson 1978). Thus, some have argued that DOL could itself be a colony level adaptation (Jeanson et al. 2007; Holbrook et al. 2011). However, the self-organization perspective argues that the scaling of DOL with group size could emerge spontaneously (Jeanson et al. 2007; Jeanson and Fewell 2008; Holbrook et al. 2013) and become a target of selection only thereafter (Page 1997; Camazine et al. 2001). In *S. dumicola*, it is unknown if DOL is functionally advantageous for colonies. However, in other species of group-living spider, enhanced DOL and greater within-group behavioral variation are both advantageous for colony mass gain (Pruitt and Riechert 2011). Thus, is seems plausible that DOL could be advantageous in *S. dumicola*, and its positive effects might increase with increasing group size. At present, much more work is needed to critically evaluate these hypotheses.

The slope of the group size vs. DOL scalar relationship in *S. dumicola* appears steeper than that those observed in some eusocial insects. We observed a nearly 10-fold increase in DOL_indiv_ across groups ranging from 6-40 spiders. This exceeds the relative increase in DOL observed among *P. californicus* harvester ant colonies ranging from 30 to 390 workers (Holbrook et al. 2011), but approximates the percent change in DOL predicted under certain simulation model conditions (Jeanson et al. 2007). When averaged across group sizes, the levels of DOL observed in *S. dumicola*, a cooperative breeder, equals or exceeds those values reported from several eusocial insects (Dornhaus et al. 2009; Jandt et al. 2009; Holbrook et al. 2011). Both of these findings together indicate that colonies of *S. dumicola* may be especially prone to the emergence of DOL. One ultimate explanation for this outcome is that the advantages of increased task specialization may be greater in these spiders than in some insects. Alternatively, the costs of task switching may be greater for social spiders than in other systems (Smith 1776). A possible proximate explanation is that the compact and labyrinthine nests of *S. dumicola* restrict or bias individuals’ opportunities to engage in some tasks versus others. More studies on *S. dumicola* and other taxa are therefore needed to discern why variation in DOL and its scalar relationship with group size should differ across systems. Such studies will enhance our understanding of why DOL emerges, adaptively or otherwise, and whether or how selection can operate on this organizational feature.

## Acknowledgements

We would like to thank the following UC Santa Barbara undergraduates for their help in the collection of the data for this project: Taylor Beverage, Kathrin Blank, Audrey Boer, Vicky Chan, Eric Clayborn, Kristina DeFalco, Colby Dunford, John Espinosa, April Fenster, Samantha Gaston, Michelle Gertsvolf, Adrene Garabedian, Briana Gibbs, Kelly Happ, Kaylyn Harris, Jake Harries, Emily Hershman, Aaron Huelsman, Matthew Jones, Emma Knutson, Aimee Miller, Nathaniel Mooi, Gabriella M. Najm, Elizabeth Orgon, Catherine J. Patton, Grant Perri, Harrison Pollard, Meghna Rao, Hayley Simon, Keenan D. Sucov, Elaine Tan, Francis Wang, Neha Yilayavilli. We would also like to thank the owners of the Tramonto Lodge in Upington, South Africa for providing us with a place to stay during our time in South Africa. Funding for this research was generously provided by NSF IOS grants 1352705, 1455895 to JNP, 1456010 376 to NPW, and NIH GM115509 to JNP and NPW.

## Literature Cited

Beshers SN, Fewell J. N. (2001) Models of division of labor in social insects. Annual Review in Entomology 46:413–440

Bonabeau E, Theraulaz G, Deneubourg JL (1996) Quantitative study of the fixed threshold model for the regulation of division of labour in insect societies. Proceedings of the Royal Society B-Biological Sciences 263:1565–1569

Bonabeau E, Theraulaz G, Deneubourg JL (1998) Fixed response thresholds and the regulation of division of labor in insect societies. Bulletin of Mathematical Biology 60:753–807

Bonner JT (2004) Perspective: The size-complexity rule. Evolution 58:1883–1890

Camazine S, Deneubourg J-L, Franks NR, Sneyd J, Theraulaz G, Bonabeau E (2001) Self-organization in biological systems. Self-organization in biological systems:i-viii, 1–538

Dornhaus A (2008) Specialization Does Not Predict Individual Efficiency in an Ant. Plos Biology 6:2368–2375

Dornhaus A, Holley J-A, Franks NR (2009) Larger colonies do not have more specialized workers in the ant Temnothorax albipennis. Behavioral Ecology 20:922–929

Duarte A, Weissing FJ, Pen I, Keller L (2011a) An Evolutionary Perspective on Self-Organized Division of Labor in Social Insects. Annual Review of Ecology, Evolution, and Systematics, Vol 42 42:91–110

Duarte A, Weissing FJ, Pen I, Keller L (2011b) An Evolutionary Perspective on Self-Organized Division of Labor in Social Insects. Annual Review of Ecology, Evolution, and Systematics 42:91–110

Foelix R (2011) Biology of spiders. Third edition. Biology of spiders Third edition:i-viii, 1–419

Gautrais J, Theraulaz G, Deneubourg JL, Anderson C (2002) Emergent polyethism as a consequence of increased colony size in insect societies. Journal of Theoretical Biology 215:363–373

Gordon DM (1989) DYNAMICS OF TASK SWITCHING IN HARVESTER ANTS. Animal Behaviour 38:194–204

Gorelick R, Bertram SM (2007) Quantifying division of labor: borrowing tools from sociology, sociobiology, information theory, landscape ecology, and biogeography. Insectes Sociaux 54:105–112

Gorelick R, Bertram SM, Killeen PR, Fewell JH (2004) Normalized mutual entropy in biology: Quantifying division of Labor. American Naturalist 164:677–682

Holbrook CT, Barden PM, Fewell JH (2011) Division of labor increases with colony size in the harvester ant Pogonomyrmex californicus. Behavioral Ecology 22:960–966

Holbrook CT, Kukuk PF, Fewell JH (2013) Increased group size promotes task specialization in a normally solitary halictine bee. Behaviour 150:1449–1466

Holbrook CT, Wright CM, Pruitt JN (2014) Individual differences in personality and behavioural plasticity facilitate division of labour in social spider colonies. Animal Behaviour 97:177–183

Holldobler B, Wilson EO (1990) The ants. Belknap Press, Cambridge, MA

Jakob EM, Marshall SD, Uetz GW (1996) Estimating fitness: A comparison of body condition indices. Oikos 77:61–67

Jandt JM, Huang E, Dornhaus A (2009) Weak specialization of workers inside a bumble bee (Bombus impatiens) nest. Behavioral Ecology and Sociobiology 63:1829–1836

Jeanne RL (1986) The organization of work in *Polybia occidentalis* - costs and benefits of specialization in a social wasp. Behavioral Ecology and Sociobiology 19:333–341

Jeanne RL (2016) Division of labor is not a process or a misleading concept. Behavioral Ecology and Sociobiology 70:1109–1112

Jeanson R, Fewell JH (2008) Influence of the social context on division of labor in ant foundress associations. Behavioral Ecology 19:567–574

Jeanson R, Fewell JH, Gorelick R, Bertram SM (2007) Emergence of increased division of labor as a function of group size. Behavioral Ecology and Sociobiology 62:289–298

Jeanson R, Kukuk PF, Fewell JH (2005) Emergence of division of labour in halictine bees: contributions of social interactions and behavioural variance. Animal Behaviour 70:1183–1193

Jeanson R, Weidenmuller, A. (2013) Interindividual variability in social insects - proximate causes and ultimate consequences. Biological Reviews

Jolles JW, Fleetwood-Wilson A, Nakayama S, Stumpe MC, Johnstone RA, Manica A (2015) The role of social attraction and its link with boldness in the collective movements of three-spined sticklebacks. Animal Behaviour 99:147–153

Junghanns A, Holm C, Schou MF, Sorensen AB, Uhl G, Bilde T (2017) Extreme allomaternal care and unequal task participation by unmated females in a cooperatively breeding spider. Animal Behaviour 132:101–107

Kay A, Rissing SW (2005) Division of foraging labor in ants can mediate demands for food and safety. Behavioral Ecology and Sociobiology 58:165–174

Keiser CN, Jones DK, Modlmeier AP, Pruitt JN (2014) Exploring the effects of individual traits and within-colony variation on task differentiation and collective behavior in a desert social spider. Behavioral Ecology and Sociobiology 68:839–850

Keiser CN, Pinter-Wollman N, Ziemba MJ, Kothamasu KS, Pruitt JN (2017) The index case is not enough: Variation among individuals, groups, and social networks modify bacterial transmission dynamics. Journal of Animal Ecology

Keiser CN, Wright CM, Pruitt JN (2016) Increased bacterial load can reduce or negate the effects of keystone individuals on group collective behaviour. Animal Behaviour 114:211–218

Koolhaas JM, Korte SM, De Boer SF, Van Der Vegt BJ, Van Reenen CG, Hopster H, De Jong IC, Ruis MAW, Blokhuis HJ (1999) Coping styles in animals: current status in behavior and stress-physiology. Neuroscience and Biobehavioral Reviews 23:925–935

Kurvers R, Eijkelenkamp B, van Oers K, van Lith B, van Wieren SE, Ydenberg RC, Prins HHT (2009) Personality differences explain leadership in barnacle geese. Animal Behaviour 78:447–453

Kurvers R, van Oers K, Nolet BA, Jonker RM, van Wieren SE, Prins HHT, Ydenberg RC (2010) Personality predicts the use of social information. Ecology Letters 13:829–837

Laskowski KL, Montiglio PO, Pruitt JN (2016) Individual and Group Performance Suffers from Social Niche Disruption. American Naturalist 187:776–785

Merkle D, Middendorf M (2004) Dynamic polyethism and competition for tasks in threshold reinforcement models of social insects. Adaptive Behavior 12:251–262

Michener CD (1974) The social behavior of the bees: a comparative study, vol 73. Harvard University Press

Oster G, Wilson EO (1978) Castes and Ecology in the Social Insects. Princeton University Press, Princeton, NJ

Oster G, Wilson, E. O. (1978) Castes and Ecology in the Social Insects. Princeton University Press, Princeton, NJ

Page RE (1997) The evolution of insect societies. Endeavour 21:114–120

Pinter-Wollman N, Mi, Brian, Pruitt JN (2017) Replacing bold individuals has a smaller impact on group performance than replacing shy individuals. Behavioral Ecology

Pruitt JN (2013) A real-time eco-evolutionary dead-end strategy is mediated by the traits of lineage progenitors and interactions with colony invaders. Ecology Letters 16:879–886

Pruitt JN, Cote J, Ferrari MCO (2012) Behavioural trait variants in a habitat-forming species dictate the nature of its interactions with and among heterospecifics. Functional Ecology 26:29–36

Pruitt JN, Keiser CN (2014) The personality types of key catalytic individuals shape colonies’ collective behaviour and success. Animal Behaviour 93:87–95

Pruitt JN, Riechert SE (2011) How within-group behavioural variation and task efficiency enhance fitness in a social group. Proceedings of the Royal Society B-Biological Sciences 278:1209–1215

Pruitt JN, Wright CM, Lichtenstein JLL, Chism GT, McEwen BL, Kamath A, Pinter-Wollman N (2017) Selection for Collective Aggressiveness Favors Social Susceptibility in Social Spiders. Current Biology 28:100–105

Riechert SE, Jones TC (2008) Phenotypic variation in the social behaviour of the spider Anelosimus studiosus along a latitudinal gradient. Animal Behaviour 75:1893–1902

Ruch J, Heinrich L, Bilde T, Schneider JM (2009) Relatedness facilitates cooperation in the subsocial spider, Stegodyphus tentoriicola. Bmc Evolutionary Biology 9

Rypstra AL (1993) PREY SIZE, SOCIAL COMPETITION, AND THE DEVELOPMENT OF REPRODUCTIVE DIVISION-OF-LABOR IN SOCIAL SPIDER GROUPS. American Naturalist 142:868–880

Seeley TD (1982) Adaptive significance of the age polyethism schedule in honeybee colonies. Behavioral Ecology and Sociobiology 11:287–293

Seibt U, Wickler W (1988) Bionomics and social structure of’Family Spiders’ of the genus Stegodyphus, with special reference to the African species S. dumicola and S. mimosarum (Araneida, Eresidae). Verhandlungen des naturwissenschaftlichen 30:255–303

Smith A (1776) The Wealth of Nations. Methuen & Co., Ltd., London

Szathmary E, Smith JM (1995) The major evolutionary transitions. Nature 374:227–232

Thomas ML, Elgar MA (2003) Colony size affects division of labour in the ponerine ant Rhytidoponera metallica. Naturwissenschaften 90:88–92

Trompf L, Brown C (2014) Personality affects learning and trade-offs between private and social information in guppies, Poecilia reticulata. Animal Behaviour 88:99–106

Ulrich Y, Saragosti J, Tokita CK, Tarnita CE, Kronauer DJC (2018) Fitness benefits and emergent division of labour at the onset of group living. Nature 560:635-+

Wilson EO (1971) The Insect Societies. Harvard University Press, Cambridge, MA

Wilson EO (1980) Caste and division of labor in leaf-cutter ants (Hymenoptera, Formicidae, Atta. The ergonomic optimization of leaf cutting. Behavioral Ecology and Sociobiology 7:157–165

Wright CM, Holbrook CT, Pruitt JN (2014) Animal personality aligns task specialization and task proficiency in a spider society. Proceedings of the National Academy of Sciences of the United States of America 111:9533–9537

Wright CM, Keiser CN, Pruitt JN (2015) Personality and morphology shape task participation, collective foraging and escape behaviour in the social spider Stegodyphus dumicola. Animal Behaviour 105:47–54

Wright CM, Keiser CN, Pruitt JN (2016) Colony personality composition alters colony-level plasticity and magnitude of defensive behaviour in a social spider. In, Animal Behaviour, pp 175–183

